## Supplementary File for "Identifying promoter sequence architectures via a chunking-based algorithm using non-negative matrix factorisation"

#### Contents

**1 Supplementary figures from experiments on simulated data**

**2 Results on Drosophila core promoters from modENCODE**

**3 seqArchR parameters choices for all organisms**

**4 Supplementary figures of results on Drosophila core promoters from [3]**

4.1 Visualisation showing curation of seqArchR result clusters for D.melanogaster . . . . . . . . . . . . .

4.2 Cluster containing non-His2B genes with His2B genes . . . . . . . . . . . . . . . . . . . . . . . . . . .

4.3 GO terms enriched for different clusters in Drosophila melanogaster development . . . . . . . . . . .

**5 Supplementary figures of results on Zebrafish core promoters from [4]**

5.1 Visualisation showing curation of seqArchR result clusters for D.rerio . . . . . . . . . . . . . . . . . .

5.2 Overlaps between promoter sequences at all stages in D.rerio . . . . . . . . . . . . . . . . . . . . . .

5.3 Top-10 GO term enrichments for clusters . . . . . . . . . . . . . . . . . . . . . . . . . . . . . . . . .

**6 Supplementary figures of results on human cell lines core promoters from ENCODE**

6.1 Visualisation showing curation of seqArchR result clusters for H.sapiens . . . . . . . . . . . . . . . . .

6.2 Comparison of cluster architectures in shorter vs longer downstream flank scenario . . . . . . . . . .

6.3 Comparison of cluster architectures in initiator knock out vs non-knock out scenario . . . . . . . . .

**References**

[1] MRC London Institute of Medical Sciences, London, UK

[2] Institute of Clinical Sciences, Faculty of Medicine, Imperial College London, Hammersmith Hospital Campus, London, UK

### 1 Supplementary figures from experiments on simulated data

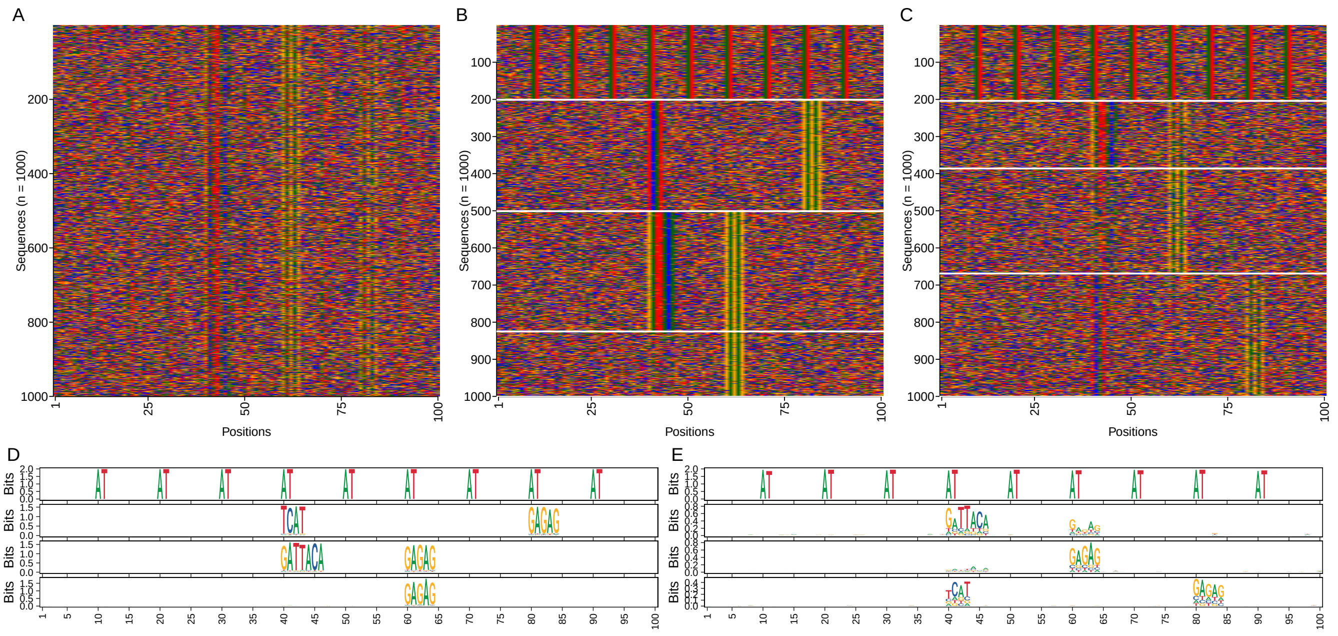


Supplementary Fig. 1: Qualitative assessment of result on simulated data. (**A**) Sequences visualised as an image for randomised input, (**B, C**) clustered output and sequence logos from seqArchR for $m = 0.1$ and $p = 1$, and (**D, E**) for $m = 0.5$ and $p = 3$.

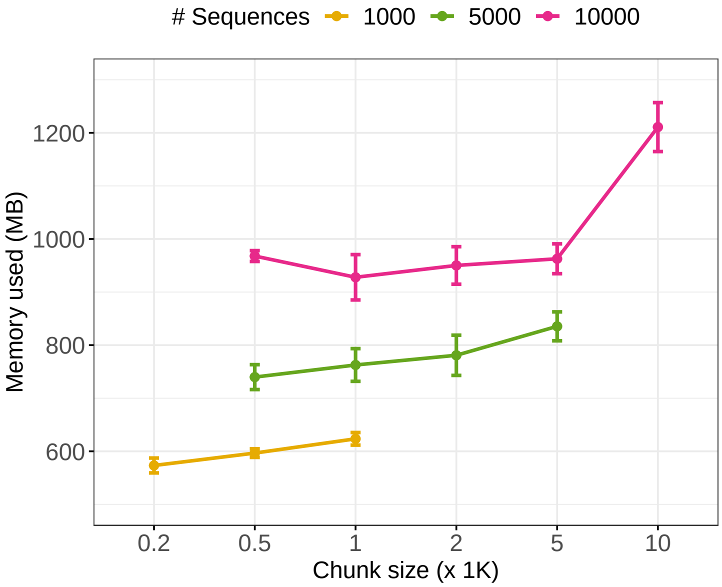


Supplementary Fig. 2: Memory footprint of seqArchR. The maximum resident set sise for different settings of seqArchR is reported.

#### 2 Results on Drosophila core promoters from modENCODE

We processed CAGE-derived core promoter sequences for Drosophila melanogaster available from modENCODE [1] with seqArchR to facilitate comparison with NPLB [2] on real promoter sequences. Specifically, 6635 core promoter sequences of D.melanogaster carcass [1] were processed with seqArchR using no parallelisation. We let seqArchR perform five iterations with collation performed at only the second iteration. While NPLB takes about 600 minutes to report 12 promoter architectures in the first pass, a comparable result from seqArchR is obtained at the end of second iteration reporting seven architectures and taking 27 minutes. NPLB further processes all the architectures identified in the first pass [ˆ1] to obtain a total of 30 architectures. [ˆ1]: Each cluster of sequences is manually processed.

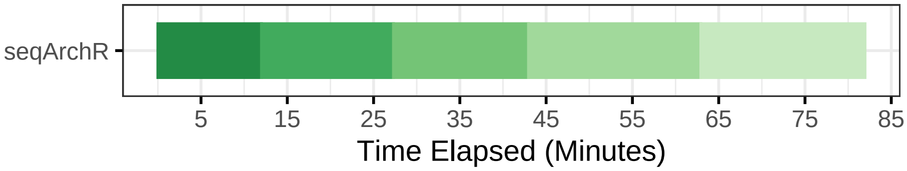


Supplementary Fig. 3: Time taken by seqArchR to process CAGE-derived core promoter sequences from D.melanogaster (modENCODE) with no parallelisation. Each shade of green represents time taken by an individual iteration – left to right/dark to light green) iteration 1 to 5.

Supplementary Figure 4 shows the architectures/clusters identified by seqArchR at the end of iteration 2. These clusters are processed further for three additional iterations. Supplementary Figure 5 shows the cluster architectures at the end of iteration 4.

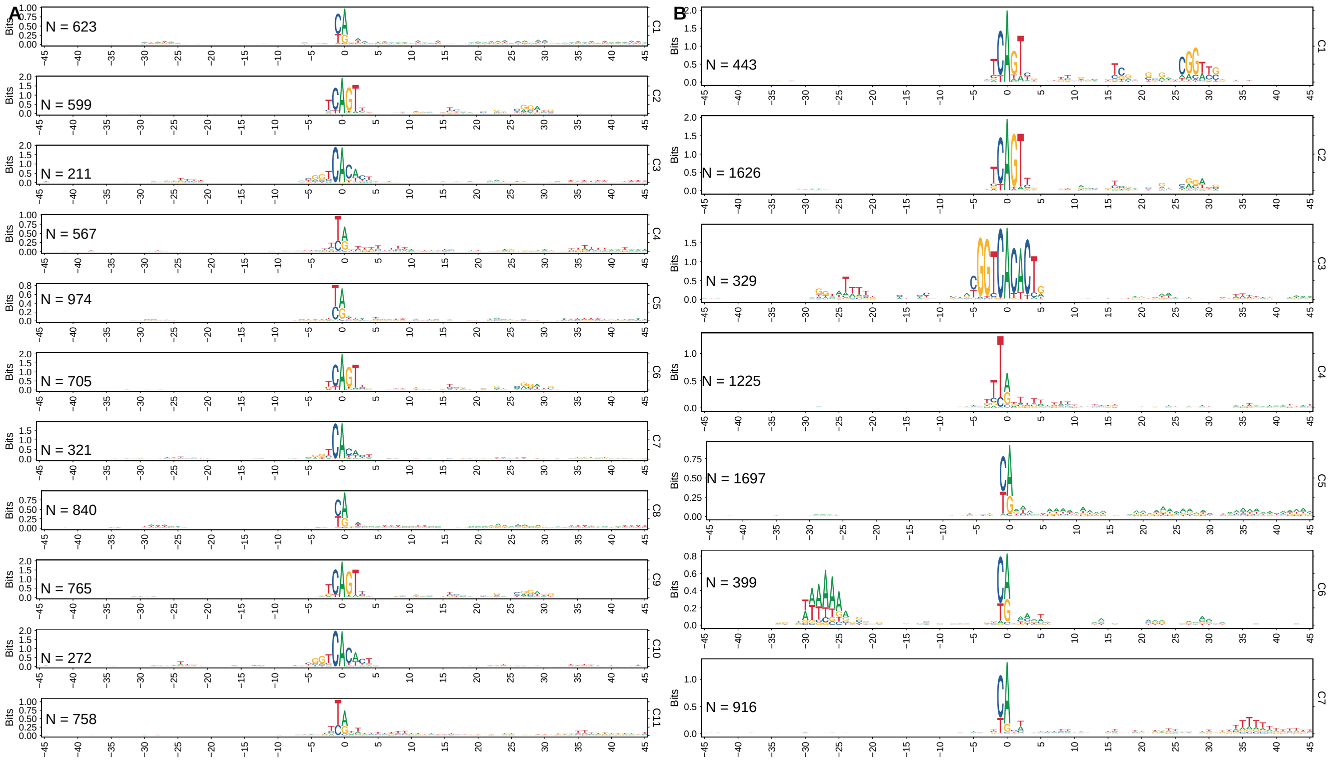


Supplementary Fig. 4: Sequence logos of promoter architectures identified by seqArchR in D. melanogaster (modENCODE) **(A)** Iteration 1; **(B)** Iteration 2.

### 3 seqArchR parameters choices for all organisms

The values chosen for different parameters for seqArchR are given in Table 1.

Table 1: Choices for each organism for preparing data and analysing with seqArchR. minTPM, minimum Tags per million, Bound value for instability of identified clusters.

| Organism | minTPM | Flank size | Bound value | #Iterations | Collation strategy |
|---|---|---|---|---|---|
| Fruit fly | 1 | −45, +45 | 1e-08 | 5 | FTTTF |
| Zebrafish | 1 | −50, +150 | 1e-06 | 5 | FTFFF |
| Humans | 1 | −50, +5 | 1e-06 | 5 | FTTTF |
| | 1 | −50, +150 | 1e-06 | 5 | FTTTF |

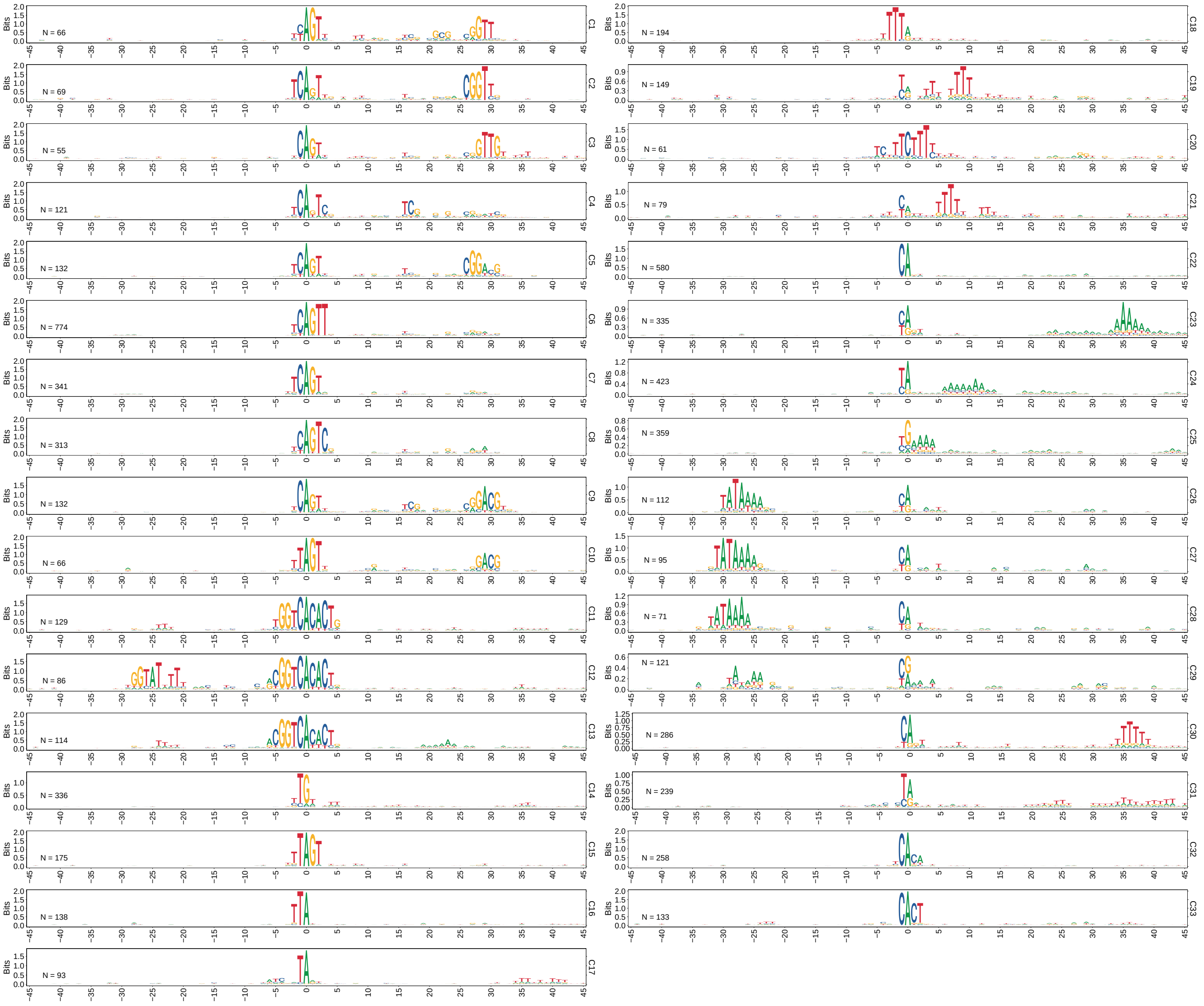


Supplementary Fig. 5: Sequence logos of promoter architectures identified by seqArchR in D. melanogaster (modENCODE) (Iteration 4)

### 4 Supplementary figures of results on Drosophila core promoters from [3]

#### 4.1 Visualisation showing curation of seqArchR result clusters for D.melanogaster

In Supplementary Figures 6, 7, and 8, we show the curation of raw clusters (of promoter sequences) from seqArchR result for the different stages processed in D. melanogaster. Note that, here, the clusters are shown ordered by the hierarchical clustering dendrogram (sequence logos on the left). Their order upon curation is arbitrary (sequence logos on the right). The final set of curated clusters is then ordered by their IQW and visualised in the figures in the main text.

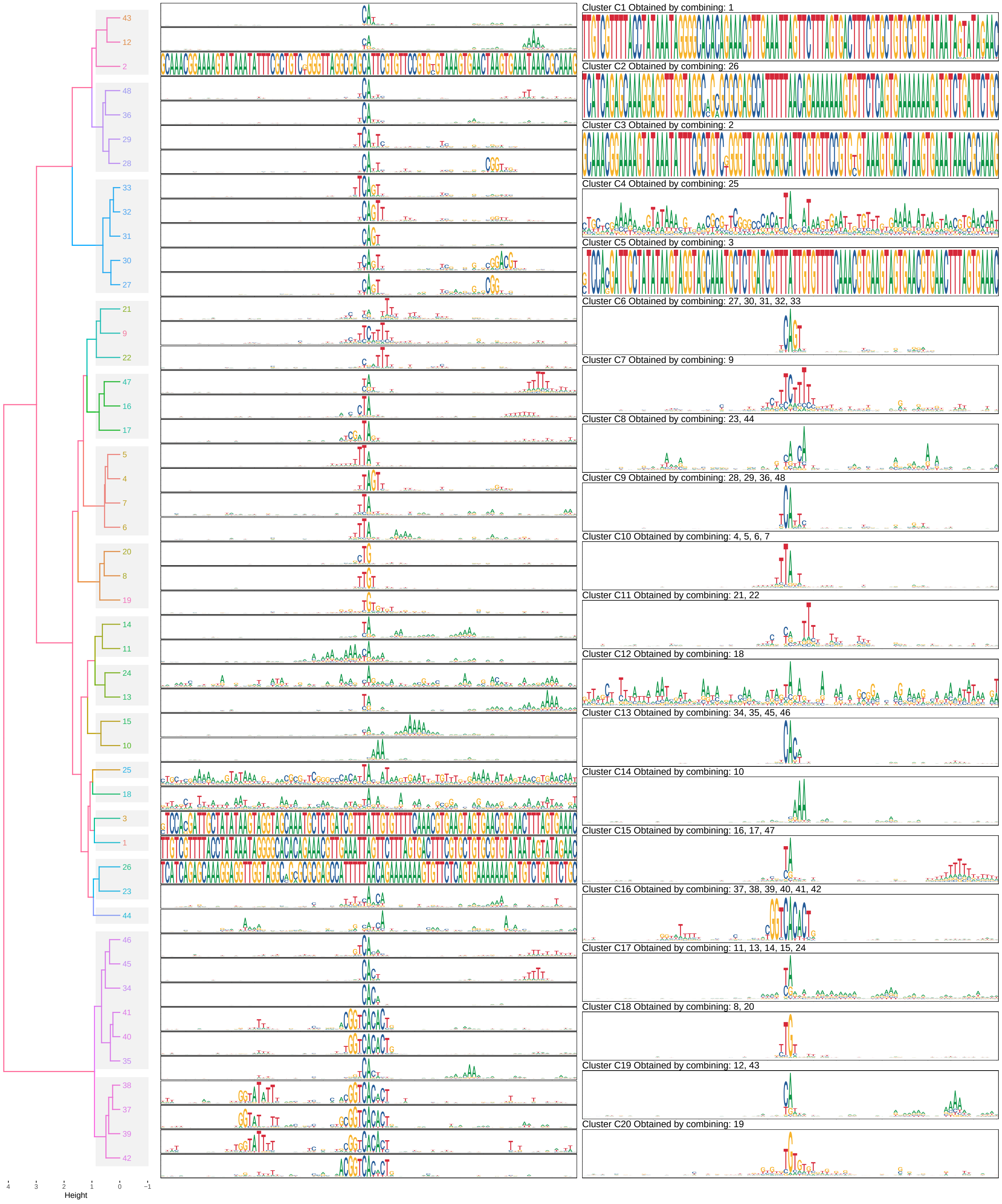


Supplementary Fig. 6: Visualisation of the collation and curation of clusters from seqArchR raw result for 2-4h AEL, D.melanogaster

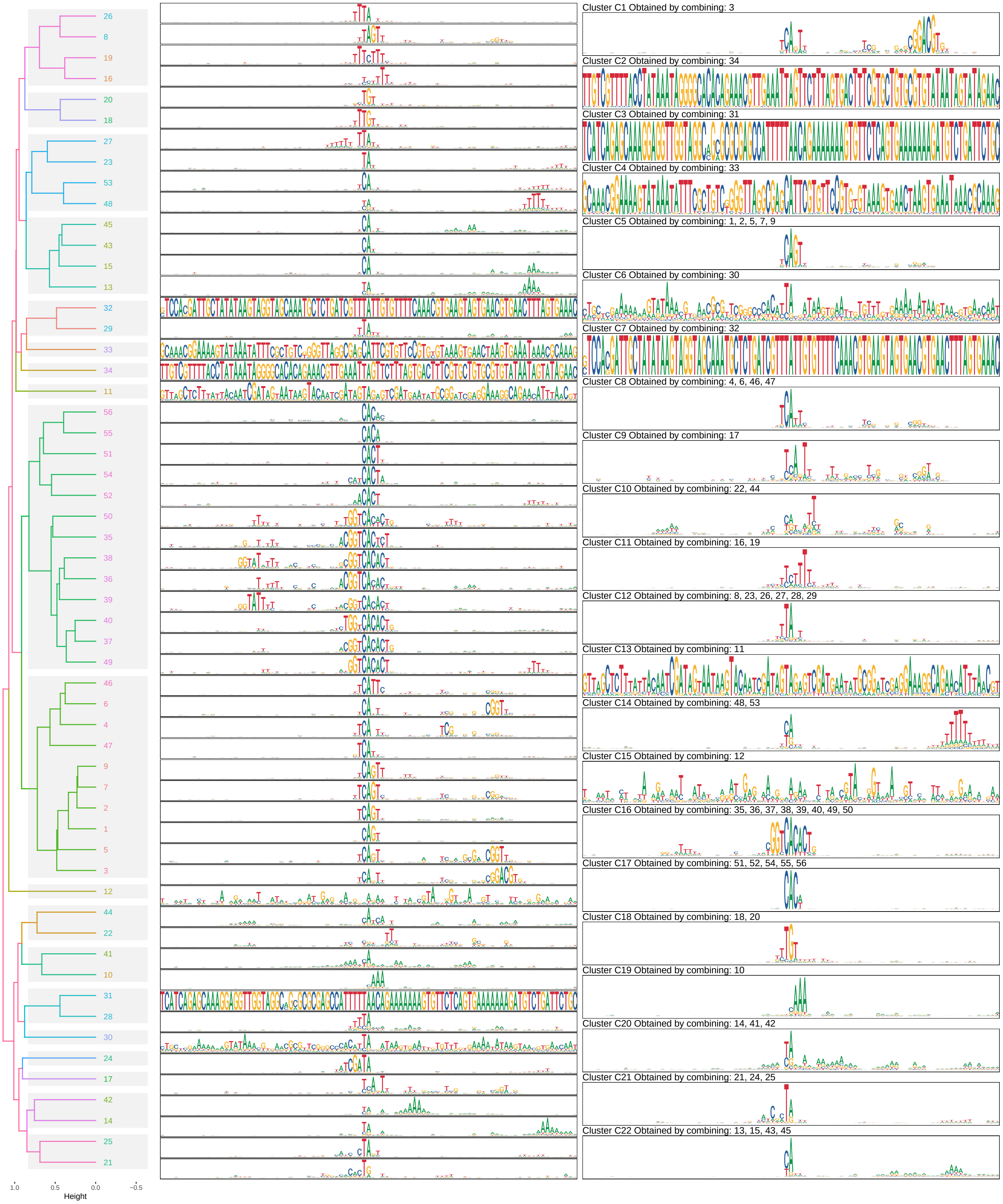


Supplementary Fig. 7: Visualisation of the collation and curation of clusters from seqArchR raw result for 6-8h AEL, D.melanogaster

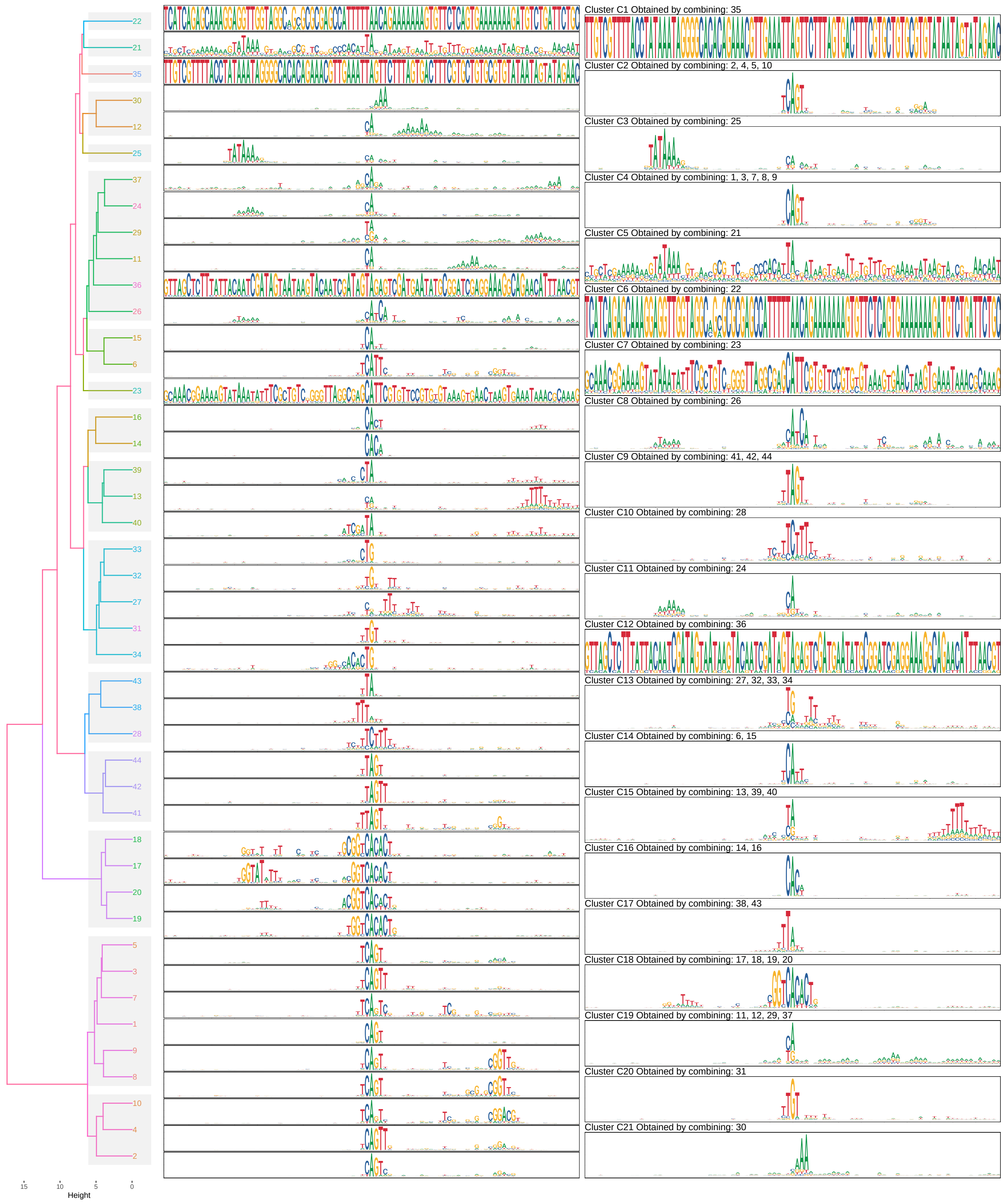


Supplementary Fig. 8: Visualisation of the collation and curation of clusters from seqArchR raw result for 10-12h AEL, D.melanogaster

##### 4.2 Cluster containing non-His2B genes with His2B genes

Observe that while most identified clusters with histone gene promoters are exclusive, only His2B promoters have been categorised with other non-His2B promoter sequences. We note that this is due to our approach to detect over-fitting (see step 2c in the Algorithm in the Methods section in the main text). NMF does separate out the non-His2B promoters from this cluster, but the over-fitting procedure adjudges it as overfit, and seqArchR, thus, re-assigns them back together with the base cluster (bringing His2B + non-His2B promoters together). Hence, the overall set of sequences remains unclustered/corrupted in the final result.

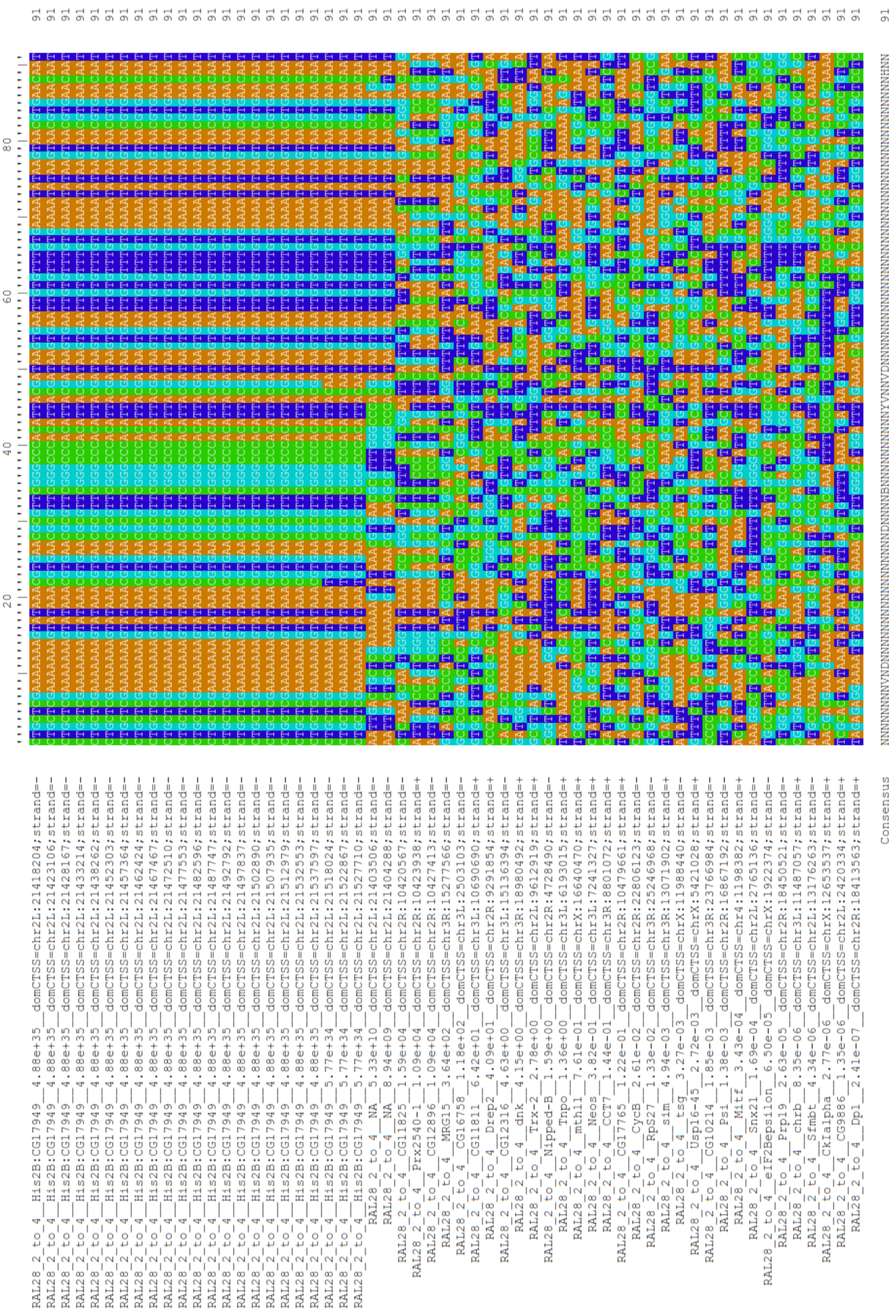

Supplementary Fig. 9: Visualisation of alignment of sequences in cluster C4X in D.melanogaster results

#### 4.3 GO terms enriched for different clusters in Drosophila melanogaster development

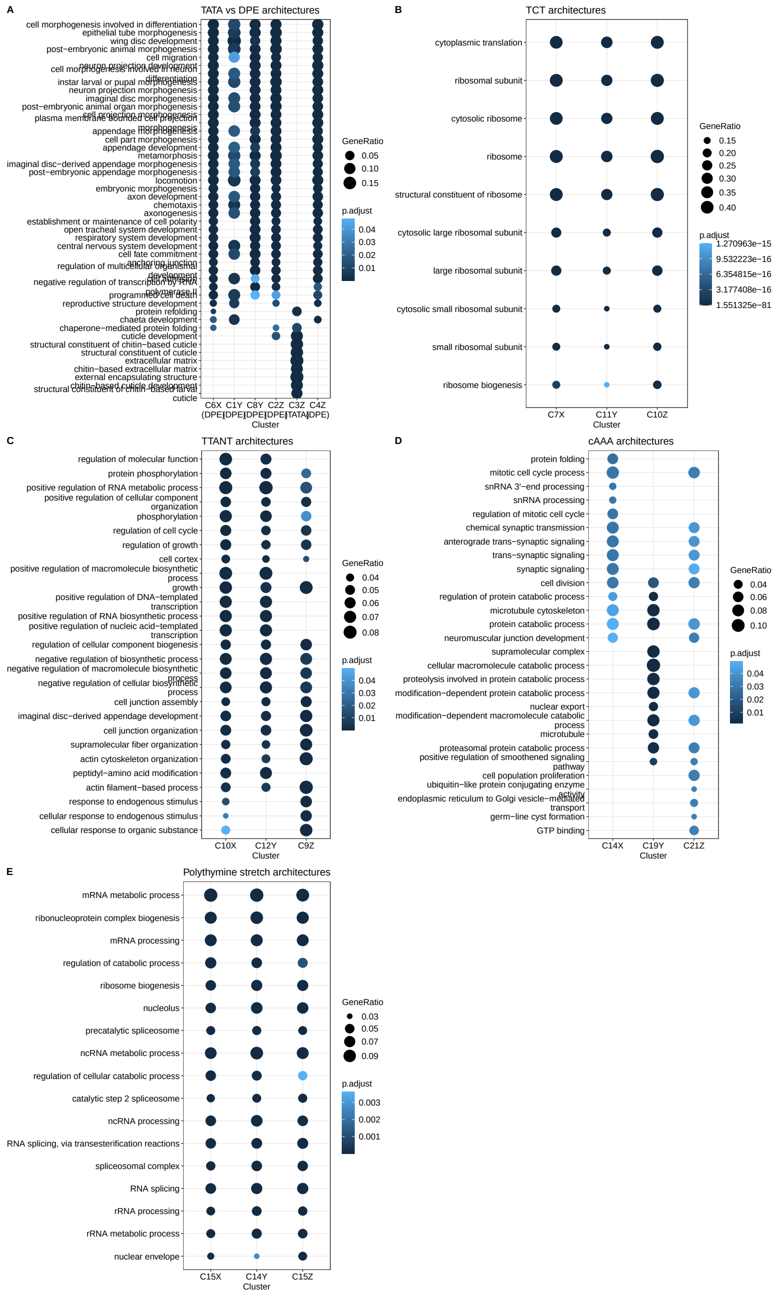


Supplementary Fig. 10: Visualisation of top-10 GO terms enriched for various clusters at different developmental stages of Drosophila melanogaster.

### 5 Supplementary figures of results on Zebrafish core promoters from [4]

#### 5.1 Visualisation showing curation of seqArchR result clusters for D.rerio

In Supplementary Figures 11, 12, and 13, we show the curation of raw clusters (of promoter sequences) from seqArchR result for the different stages processed for Danio rerio. Note that, here, the clusters are shown ordered by the hierarchical clustering dendrogram (sequence logos on the left). Their order upon curation is arbitrary (sequence logos on the right). The final set of curated clusters is then ordered by their IQW and visualised in the figures in the main text.

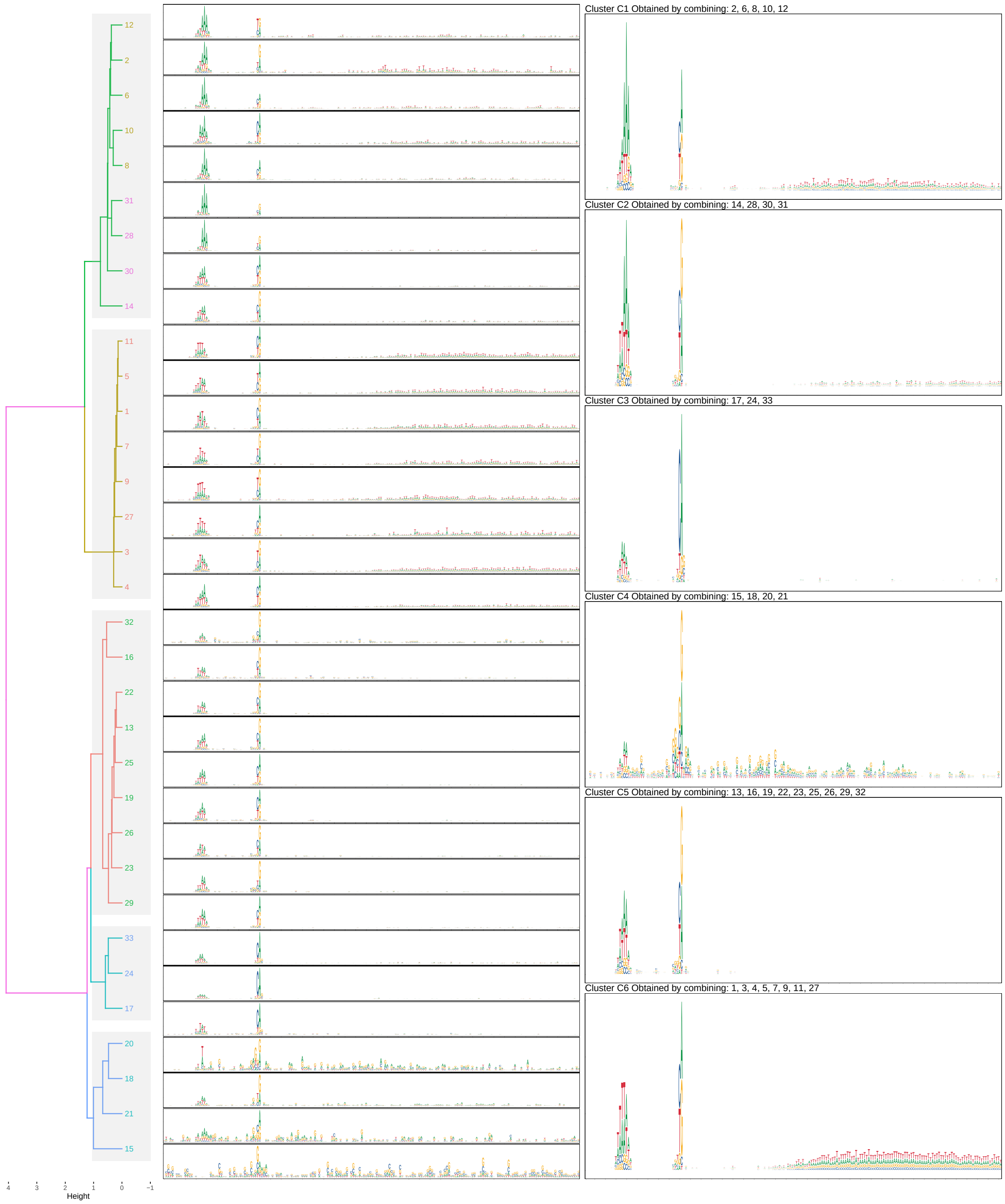


Supplementary Fig. 11: Visualisation of the collation and curation of clusters from seqArchR raw result for 64 cells stage, D.rerio

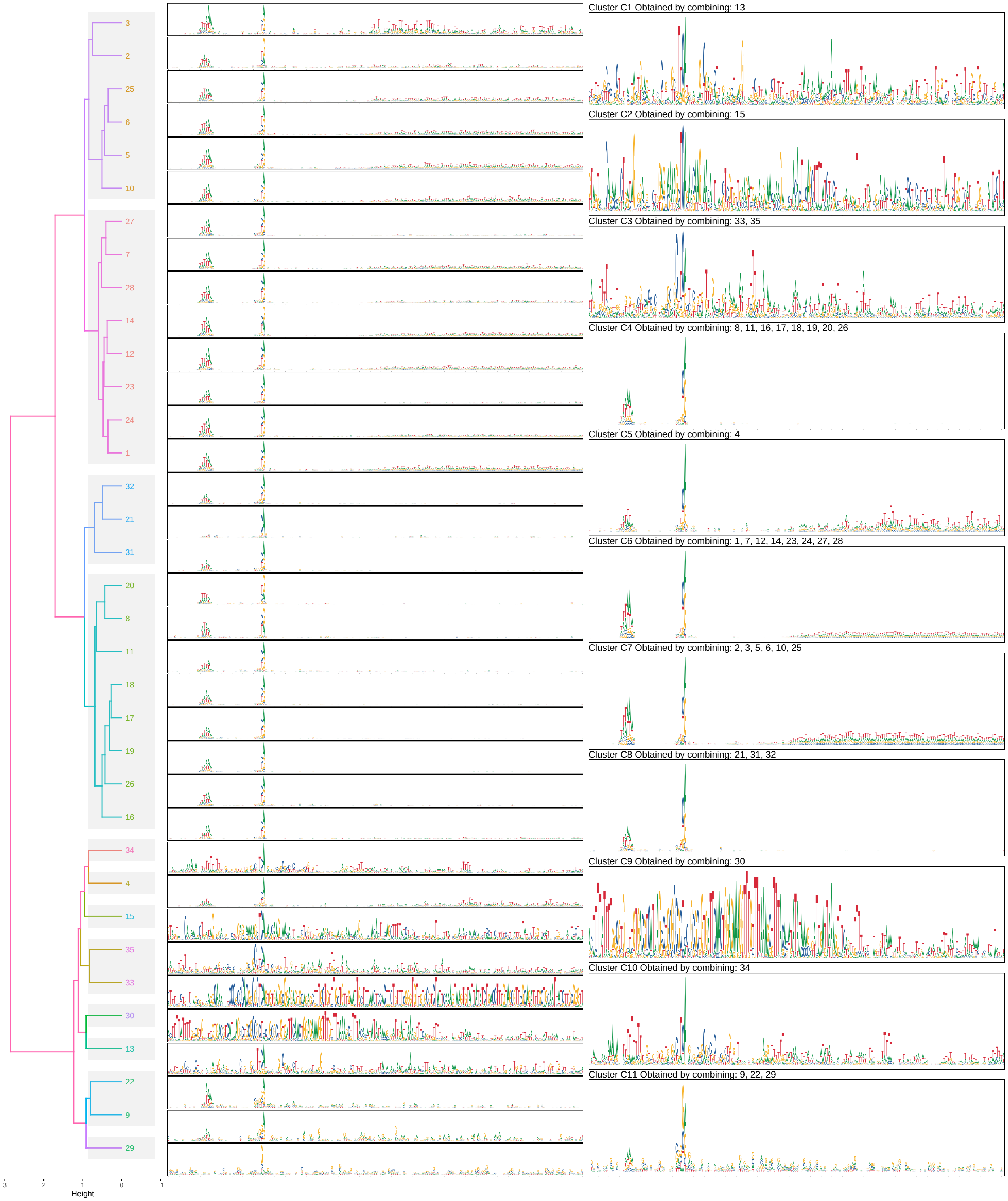


Supplementary Fig. 12: Visualisation of the collation and curation of clusters from seqArchR raw result for 30% Epiboly/Dome stage, D.rerio

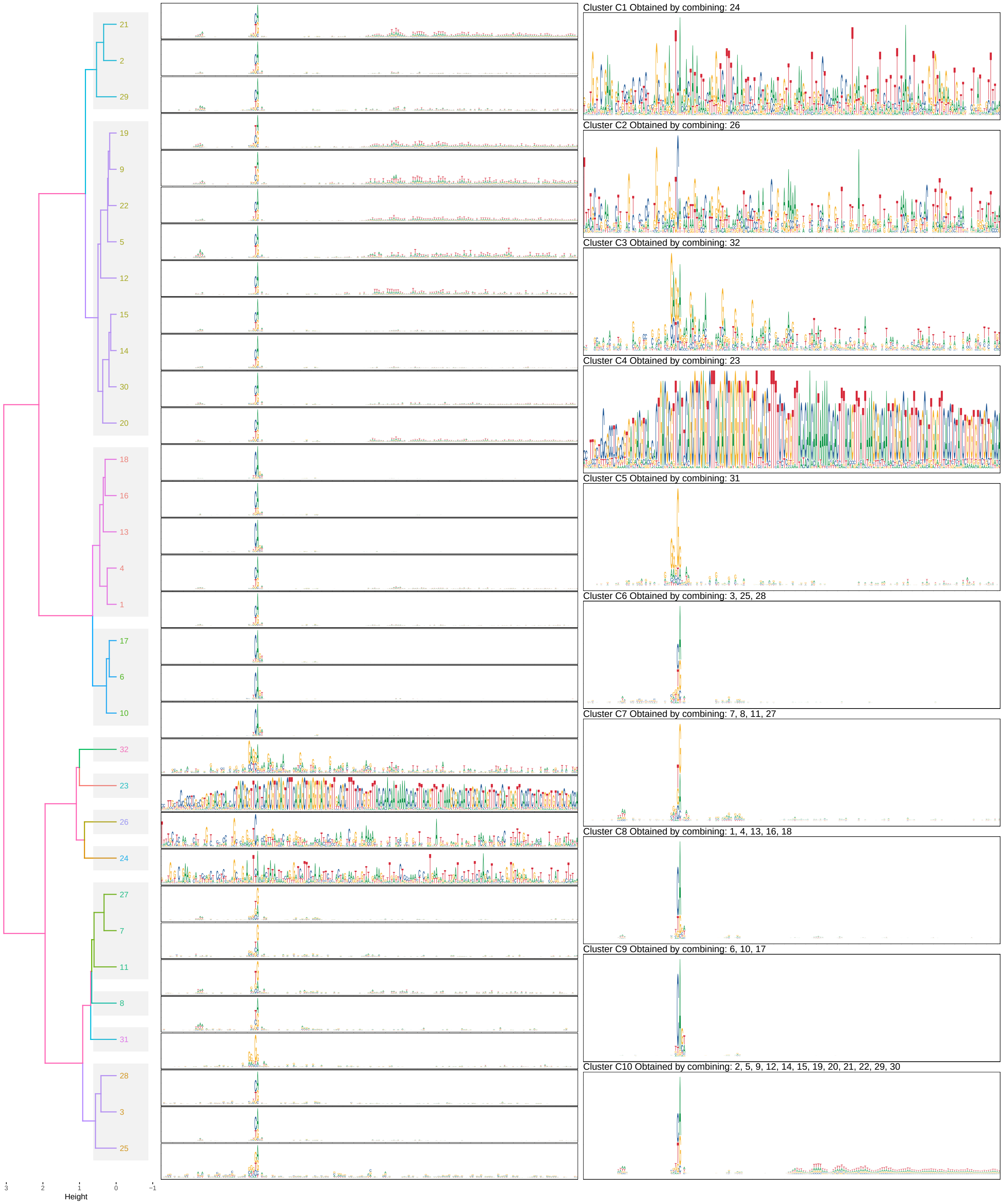


Supplementary Fig. 13: Visualisation of the collation and curation of clusters from seqArchR raw result for Prim-6, D.rerio

#### 5.2 Overlaps between promoter sequences at all stages in D.rerio

Similar to the rightmost panel figure for Drosophila stage figures, we visualise the overlaps between tag clusters at every developmental stage in zebrafish.

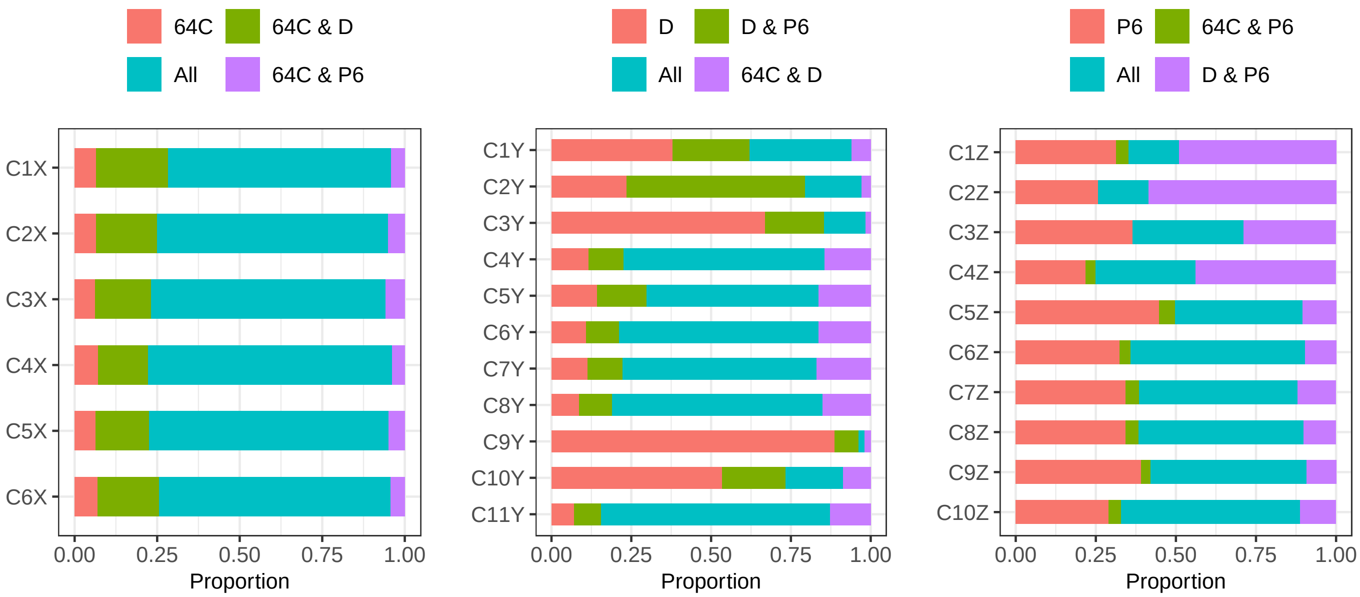


Supplementary Fig. 14: Visualisation of the overlaps between promoter sequences per developmental stage for D.rerio. Stages are abbreviated as follows. 64C: 64 cells stage; D: 30% Epiboly/Dome stage; P6: Prim-6 stage.

##### 5.3 Top-10 GO term enrichments for clusters

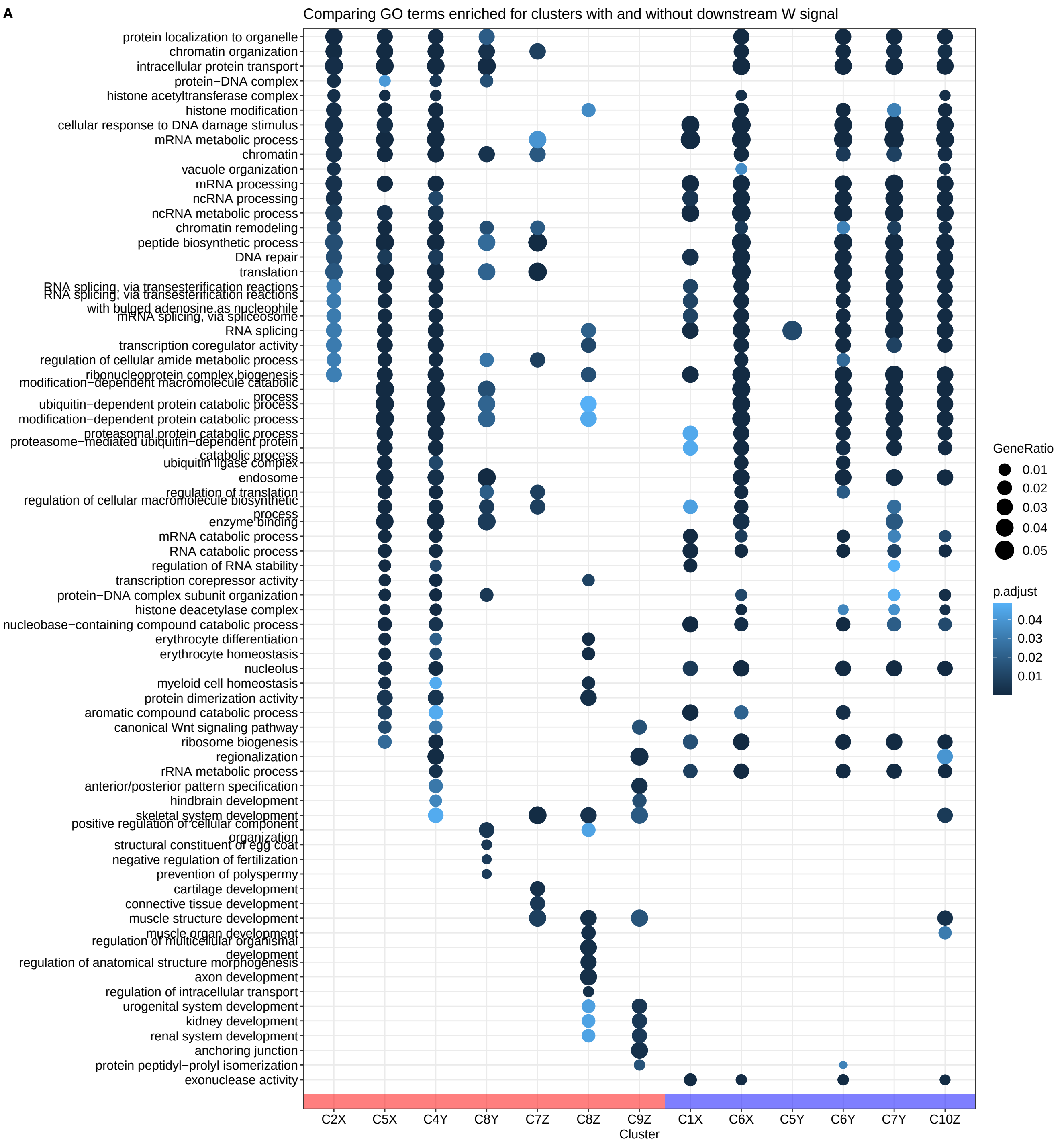


Supplementary Fig. 15: Top-10 enriched GO terms for selected clusters across ZF developmental stages.

### 6 Supplementary figures of results on human cell lines core promoters from ENCODE

We processed promoter sequences obtained using ENCODE CAGE data in human cell lines and tissues. Supplementary Figure 16 shows the library sizes of various cell lines (including replicates) obtained from ENCODE and used upon merging together. The experiments were performed with different sizes of flanks around the TSS. Specifically, upstream flank of 50 bp was combined with a short, 5 bp flank and a longer, 150 bp, flank downstream. The rationale for this is explained in the main text, the Results subsection on Homo sapiens.

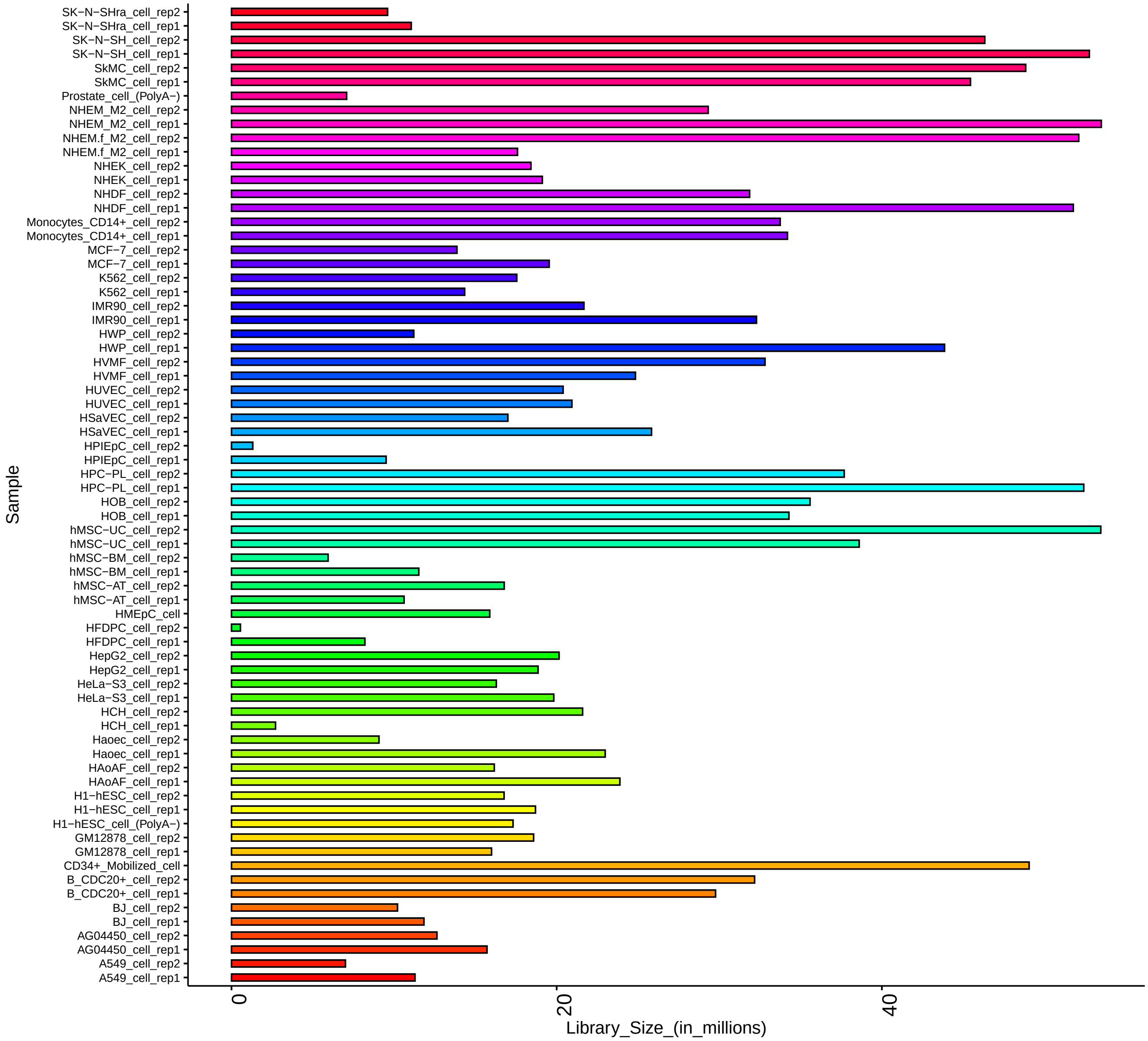


Supplementary Fig. 16: CAGE library sizes for all different human cell lines available from ENCODE and included in this study.

#### 6.1 Visualisation showing curation of seqArchR result clusters for H.sapiens

In Supplementary Figures 17, we show the curation of raw clusters (of promoter sequences) from seqArchR result for the different stages processed for Danio rerio. Note that, here, the clusters are shown ordered by the hierarchical clustering dendrogram (sequence logos on the left). Their order upon curation is arbitrary (sequence logos on the right). The final set of curated clusters is then ordered by their IQW and visualised in the figures in the main text.

#### 6.2 Comparison of cluster architectures in shorter vs longer downstream flank scenario

To observe the effect of the downstream flanking region on the clusters identified by seqArchR, we processed two sets of core promoter sequences: one with shorter downstream flank ($\{-50, +5\}$ bp) around the dominant CTSS and another with longer downstream flank ($\{-50, +150\}$ bp). Supplementary Figure 18 visualises the sequence logos of promoter architectures per cluster for both cases, shorter flanks on the left, longer flanks on the right. These are separated by a Sankey diagram that shows the movement/exchange of promoter sequences between the two sets of clusters.

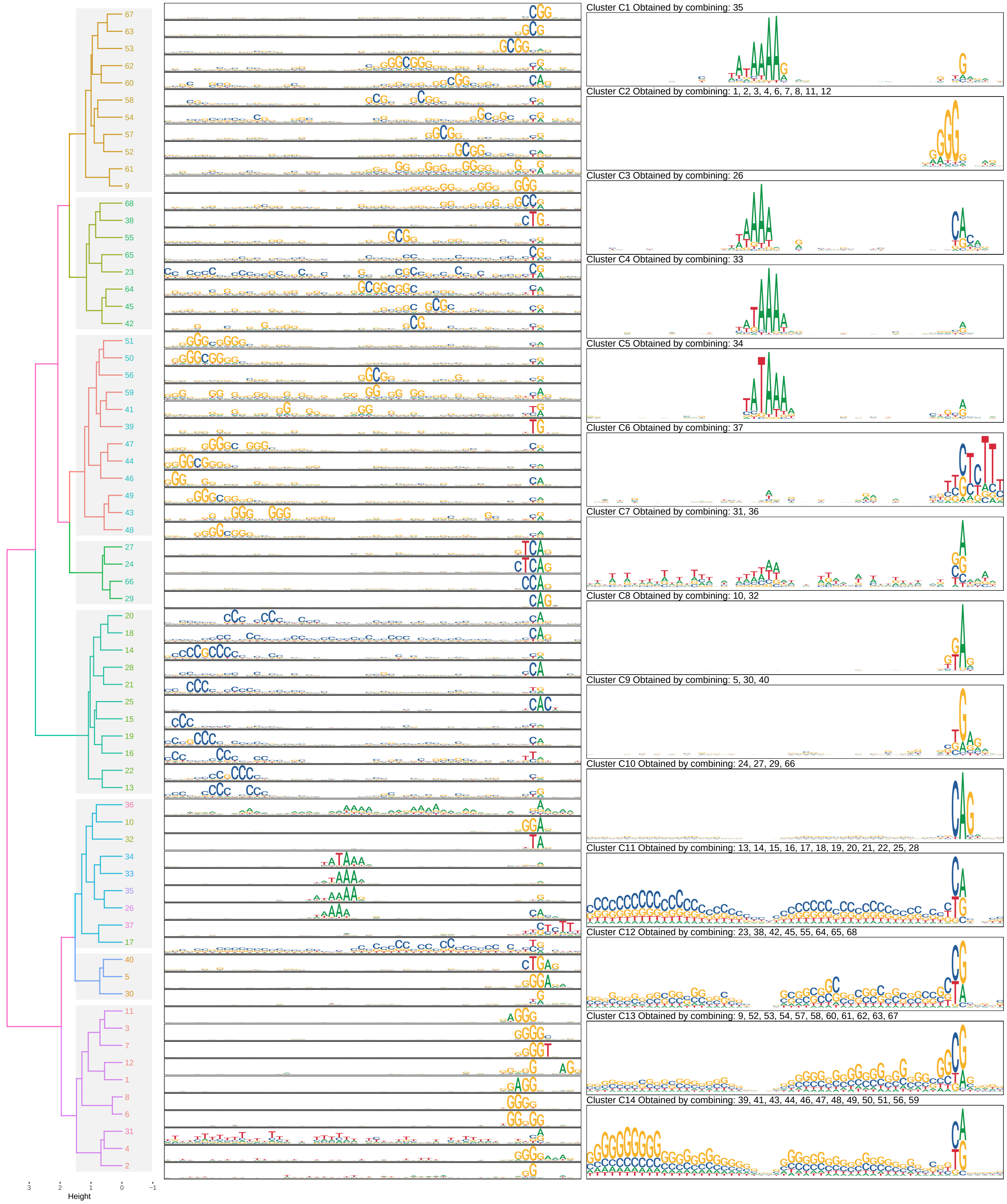


Supplementary Fig. 17: Visualisation of the collation and curation of clusters from seqArchR raw result for 2-4h AEL, D.melanogaster

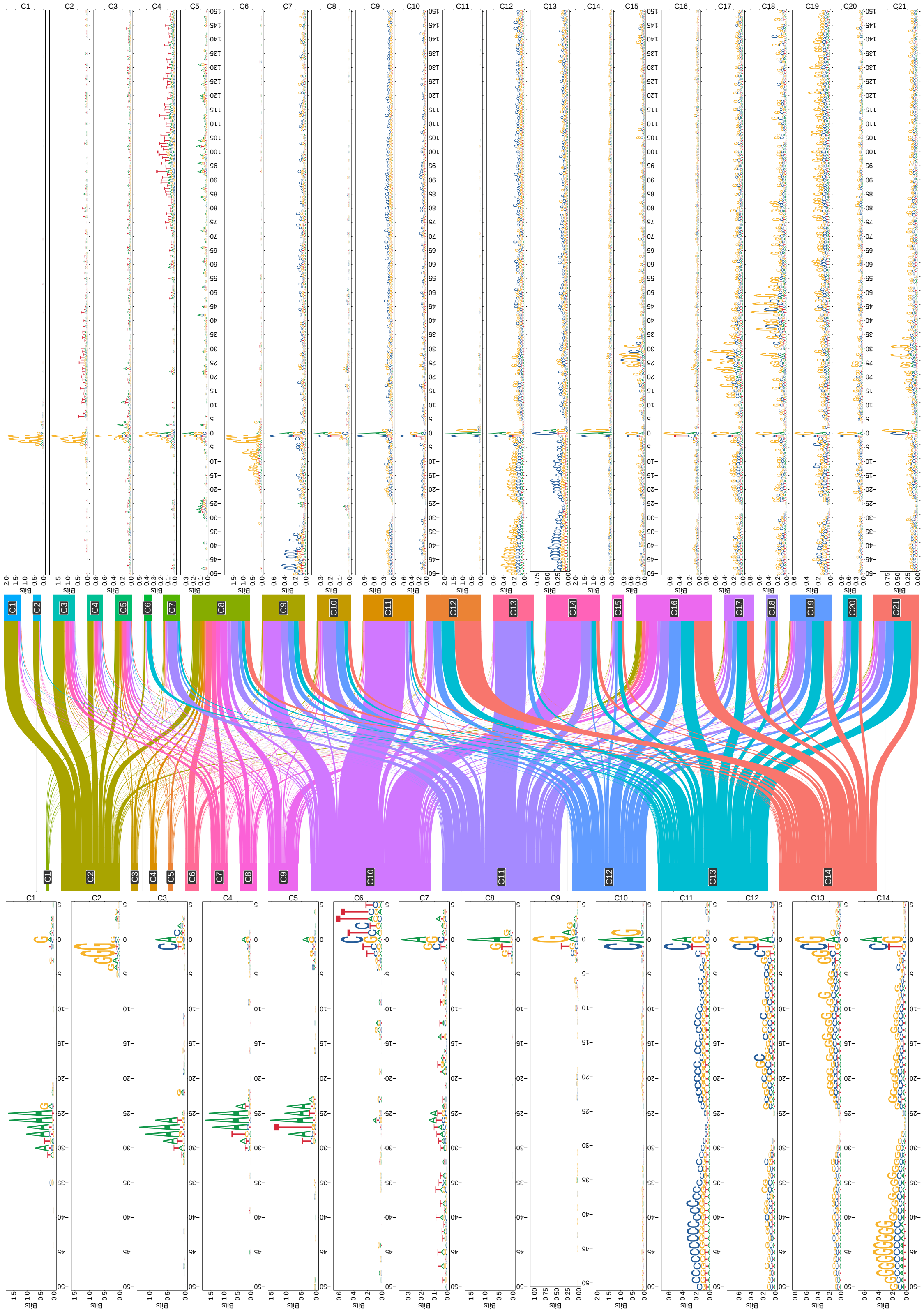


Supplementary Fig. 18: Sankey diagram showing movement of promoter sequences in two clusterings: using flanks of {−50, +5} and {−50, +150} around the dominant TSS.

#### 6.3 Comparison of cluster architectures in initiator knock out vs non-knock out scenario

To observe the effect of the initiator sequence (Inr) on the clustering, we excluded the Inr sequence positions from clustering and compared how the clustering of the core promoter sequences changed. This is visualised in Supplementary Figure 19. The sequence logos on the left are for clusters obtained by retaining/using the Inr sequence $[-5, +5$ around the TSS$]$ when clustering vs excluding it when performing clustering on the right. The Sankey diagram in the middle shows the movement of promoter sequences in the two clusterings. Note that the position labels for the sequence logos on the right go from $\{-50, -6\}$ and $\{+6, +150\}$.

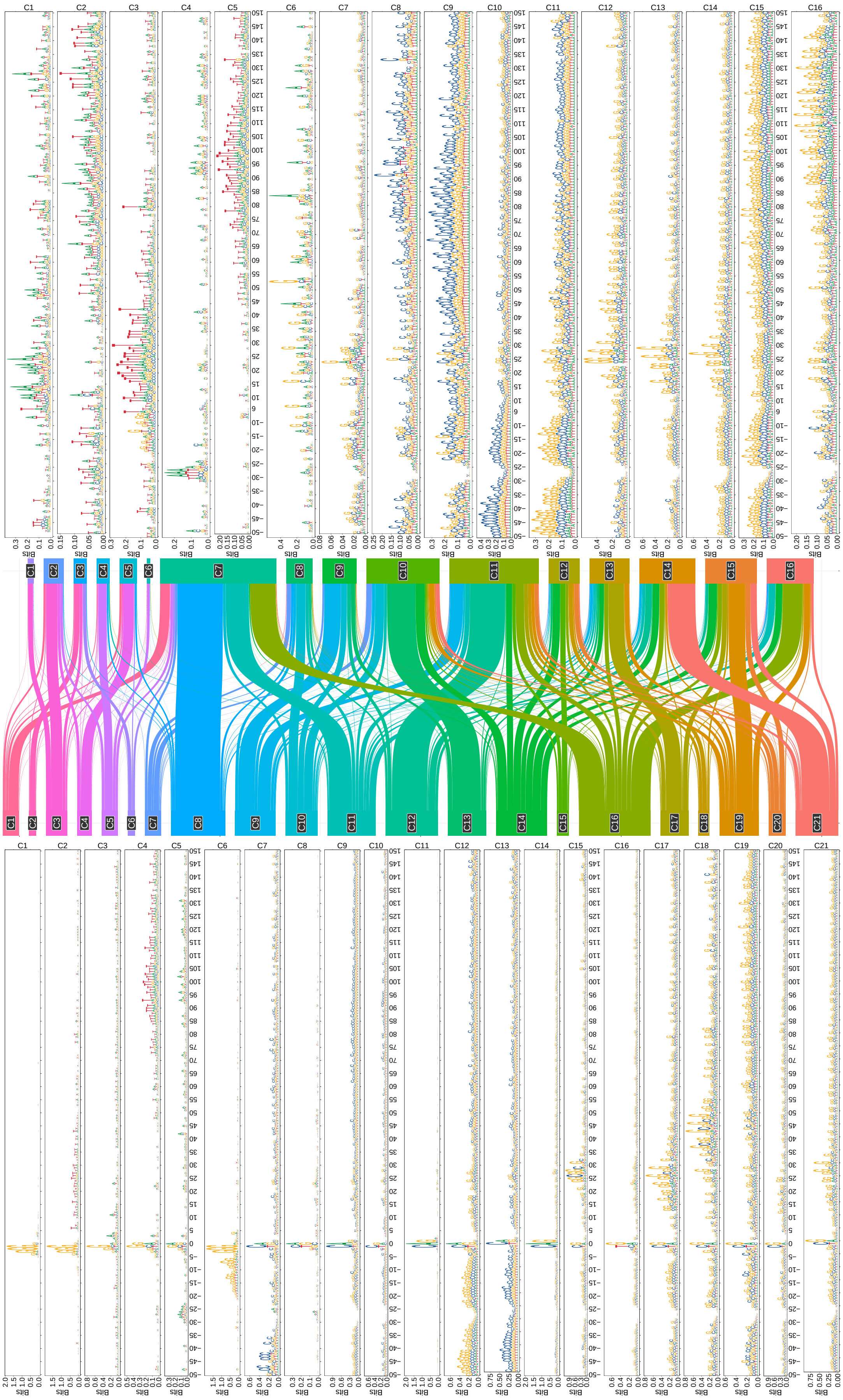


Supplementary Fig. 19: Sankey diagram showing effect of Inr knockout on the clustering.
